# SNP genotyping and parameter estimation in polyploids using low-coverage sequencing data

**DOI:** 10.1101/120261

**Authors:** Paul D. Blischak, Laura S. Kubatko, Andrea D. Wolfe

**Affiliations:** Dept. of Evolution, Ecology, and Organismal Biology; Dept. of Statistics, The Ohio State University, Columbus, OH 43210, USA.

## Abstract

**Motivation:** Genotyping and parameter estimation using high throughput sequencing data are everyday tasks for population geneticists, but methods developed for diploids are typically not applicable to polyploid taxa. This is due to their duplicated chromosomes, as well as the complex patterns of allelic exchange that often accompany whole genome duplication (WGD) events. For WGDs within a single lineage (auto polyploids), inbreeding can result from mixed mating and/or double reduction. For WGDs that involve hybridization (allopolyploids), alleles are typically inherited through independently segregating subgenomes.

**Results:** We present two new models for estimating genotypes and population genetic parameters from genotype likelihoods for auto- and allopolyploids. We then use simulations to compare these models to existing approaches at varying depths of sequencing coverage and ploidy levels. These simulations show that our models typically have lower levels of estimation error for genotype and parameter estimates, especially when sequencing coverage is low. Finally, we also apply these models to two empirical data sets from the literature. Overall, we show that the use of genotype likelihoods to model non-standard inheritance patterns is a promising approach for conducting population genomic inferences in polyploids.

**Availability:** A C++ program, EBG, is provided to perform inference using the models we describe. It is available under the GNU GPLv3 on GitHub:

https://github.com/pblischak/polyploid-genotyping.

**Contact**.

## Introduction

The discovery and analysis of genetic variation in natural populations is a central task of evolutionary genetics, with applications ranging from the inference of population structure and patterns of historical demography, detecting selection and local adaptation, and performing genetic association studies. The ability to use high throughput sequencing technologies to detect variants across the genome has further advanced our understanding of the impact of evolutionary forces on genetic diversity in populations. However, the nature of data sets collected using high throughput sequencing often require special considerations regarding sequencing error and, especially, the level of sequencing coverage. Common approaches for dealing with low-coverage sequence data use genotype likelihoods to integrate over the uncertainty of inferring genotypes when estimating other parameters [allele frequencies, inbreeding coefficients, population differentiation, etc.] (e.g., Martin *et al*., 2010; Li, 2011; Nielsen *et al*., 2011, 2012; Fumagalli *et al*., 2013; Vieira *et al*., 2013; Huang *et al*., 2016, among others). Genotype likelihoods for biallelic SNPs are calculated as the probability of the sequencing read data mapping to a variable site (total number of reads, number of reads with the alternative allele, and probability of sequencing error) given the possible values of the genotypes (typically 0,1, or 2 for the number of copies of the alternative allele in diploids). When combined with computationally efficient algorithms for inference, these models are the primary tools used for conducting population genetic analyses from high throughput data.

Although the theory for these models is well established for diploids and even special cases of higher ploidy samples (treated equivalently to pooled samples of multiple diploids), the application of these tools to taxa that have experienced a recent whole genome duplication (WGD) is currently limited (McKenna *et al*., 2010; DePristo *et al*., 2011; Li, 2011). This is due in part because of ambiguity in the copy number of each allele in the genotype of a polyploid, a phenomenon referred to as allelic dosage uncertainty (Blischak *et al*., 2016). Another important aspect of polyploid evolution to consider is that the occurrence of WGD can have an impact on how alleles are exchanged in a population, making the assumption of randomly inherited alleles inappropriate. Together these two factors have limited the widespread application of population genomic tools to gain insights about levels of genetic variation following WGD. Given both the evolutionary and economic importance of many of these organisms (e.g., agricultural crops, farmed fishes), the development of methods that can accommodate more complex patterns of inheritance is critical for the study of polyploids (Stebbins, 1950; Grant, 1971; Otto and Whitton, 2000; Soltis and Soltis, 2000; Soltis *et al*., 2014).

In this paper we present two new models for SNP genotyping in polyploids using high throughput sequencing data. The models correspond to two different ways in which polyploids can be formed: WGD within a lineage (autopolyploid) or involving hybridization between two lineages (allopolyploid). The former builds off of previous work to relax the assumption of Hardy-Weinberg equilibrium by including inbreeding (Blischak *et al*., 2016) and the latter provides a framework for separately determining the genotypes within the two genomes that compose the allopolyploid (typically referred to as subgenomes). We test our models using a wide range of simulations and describe our numerical approach for parameter estimation using the expectation maximization (EM) algorithm (Dempster *et al*., 1977). For comparison, we analyzed our simulated data sets using two additional approaches based on models that assume either Hardy Weinberg equilibrium or equal genotype probabilities. Finally, we also test the models on empirical data sets collected for a diploid-allotetraploid species pair from the genus *Betula* (birch trees) and a mixed-ploidy grass species, *Andropogon gerardii*. Overall, we demonstrate that genotype uncertainty resulting from both low-coverage sequencing data, allelic dosage uncertainty, and non-standard inheritance patterns can be overcome in polyploids using genotype likelihoods.

## Models

**Assumptions**: For each of the models below, we assume that SNPs are biallelic, and that loci and individuals are independent. For the autopolyploid model, we do not directly include double reduction (but see **Discussion**). For the allopolyploid model, we assume that subgenomes are independent, that they do not interact during meiosis (i.e., no homoeologous recombination), and that they are both in Hardy Weinberg equilibrium.

Notation for each model is introduced in the descriptions we provide below and is also summarized in Table 1. Throughout the paper, we use boldface letters to denote an array of the respective parameter across either individuals (*N*), loci (*L*), or both (e.g., ***p***:= *p*_1_, *…, p_L_*, ***F***:= *F*_1_, *…, F_N_*, and ***G***:= *g*_11_, *g*_12_, *…, g_N_* _(*L−*1)_, *g_N L_*).

**Table 1.** A key to the symbols and notation that are used in describing the autopolyploid and allopolyploid models. We use a either abold or bold-capitalized letter when referring to the collection of parameters together (e.g., ***G*** refers to *g_iℓ_* for all individuals at all loci). Parameters within subgenomes for the allopolyploid model use the same symbol but with either a1 or a2 added as a subscript.

| **Symbol** | **Description** |
| --- | --- |
| $N, L$ | The number of individuals and loci sampled. |
| $m_i$ | Ploidy level of individual $i$. |
| $d_{i\ell}$ | Sequence data for individual $i$ at locus $\ell$<br>($=\{t_{i\ell}, r_{i\ell}, \epsilon_\ell\}$). |
| $t_{i\ell}$ | Total number of reads for individual $i$ at locus $\ell$. |
| $r_{i\ell}$ | Number of alternative allele reads for individual $i$ at locus $\ell$. |
| $\epsilon_\ell$ | Average sequencing error at locus $\ell$. |
| $g_{i\ell}$ | Genotype for individual $i$ at locus $\ell$. |
| $p_\ell$ | Allele frequency at locus $\ell$. |
| $F_i$ | Inbreeding coefficient for individual $i$. |

### Autopolyploid Model

The genotype for a biallelic SNP in an autopolyploid with *K* sets of chromosomes has *K* + 1 possible values. For example, using *A* and *a* to denote the two alleles, an autotetraploid can have genotypes equal to *AAAA*, *AAAa*, *AAaa*, *Aaaa*, or *aaaa* (e.g., *g_iℓ_*= 0, 1, 2, 3, or 4, if *a* is the alternative allele; *i* = 1, *…, N* and *ℓ*= 1, *…, L*). A simple extension of the typical binomial sampling (Hardy Weinberg; HW) model used for diploids but with larger sample size to accommodate higher ploidy levels has been used previously (Li, 2011; Blischak *et al*., 2016). However, inbreeding in various forms can bias inferences made when HW equilibrium is assumed. Vieira *et al*. (2013) introduced a genotype prior to include inbreeding either per-site or per-individual for a sample of diploids (implemented in the programs ngsF and ANGSD). This model used a formulation for generalized HW that includes the inbreeding coefficient, *F*, which is the probability that two alleles are identical by decent (ibd). Instead of using a generalized HW formulation for autopolyploids, we used the Balding-Nichols beta-binomial model (Balding and Nichols, 1995, 1997; Bradburd *et al*., 2013), which also models the probability of two alleles being ibd but is more easily extended to higher ploidy levels by not directly enumerating all combinations of allele draws for the genotype of an autopolyploid. The beta-binomial distribution is obtained from the product of a binomial and beta distribution, which are commonly used in population genetics to model genotypes and allele frequencies, respectively (Wright, 1931). The beta distribution in this case is used to model genetic correlations that can result from inbreeding and/or population subdivision. These types of models are commonly referred to as *F-models* because of their relation to Wright’s fixation indices (e.g., *F_IS_*, *F_ST_*; Wright, 1931), and they form the basis of many well-known population genetic models, including those by Holsinger *et al*. (2002), Falush *et al*. (2003), and Foll and Gaggiotti (2008), as well as more recent modeling applications that include uncertainty in genotype calling from high throughput sequencing data using genotype likelihoods (e.g., Gompert *et al*., 2010; Gompert and Buerkle, 2011; Fumagalli *et al*., 2013).

Given genotype values at *L* loci for *N* individuals each of ploidy *m_i_*, we model individual genotypes at each locus (*g_iℓ_* = 0, *…, m_i_* copies of the alternative allele) as a beta-binomial random variable. This distribution derives from treating the probability of drawing an alternative allele as a beta distributed random variable with parameters 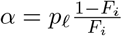 and 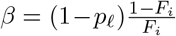, which scales the binomial probability of successfully drawing an alternative allele by both the allele frequency (*p_ℓ_*) and the amount of inbreeding (*F_i_*) (Balding and Nichols, 1995; Bradburd *et al*., 2013). The log likelihood of the genotype data for this model given the allele frequency at each site (*p_.€_*) and the per-individual inbreeding coefficients (*F_i_*) is then

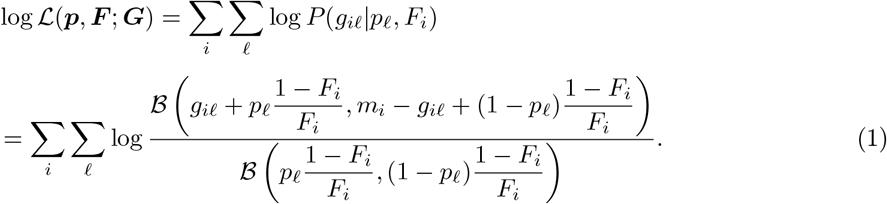

where *B*(*α, β*) represents the beta function with parameters *α* and *β*. Since genotypes must be inferred from sequence data (*d_iℓ_*; see **Methods**), we can also account for this uncertainty by summing over the possible genotype values to get the likelihood of the sequence data given allele frequencies and inbreeding coefficients by including genotype likelihoods [*P* (*d_iℓ_|g_iℓ_*)]:

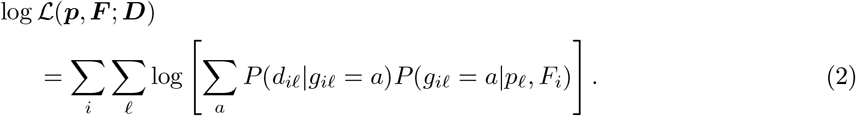

Here *P* (*g_iℓ_|p_ℓ_, F_i_*) is the beta-binomial distribution from Eq. (1). Because maximization of the log likelihood is encumbered by the logarithm of the sum over genotypes, we instead use an expectation conditional maximization algorithm to obtain maximum likelihood (ML) estimates for ***p*** and ***F*** (Meng and Rubin, 1993). Since an analytical solution for the maximization step is not readily available, we instead employ numerical maximization of the likelihood using Brent’s method (Brent, 1973). Then, given the ML parameter estimates, we can calculate the posterior probability of the genotype of each individual at each locus using Bayes’ theorem:

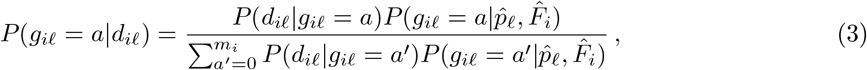

for *a* = 0, *…, m_i_*.

### Allopolyploid Model

Deviations from simple HW expectations are evident in allopolyploids in that they have two (sometimes more) sets of chromosomes inherited from separate evolutionary lineages. When these sets of chromosomes (called *homoeologs*, or *homoeologous chromosomes*) segregate during meiosis, they are inherited separately from one another and should be treated independently. For example, the genotypes for a biallelic SNP in an allotetraploid with two diploid subgenomes could have values *AA|A′A′*, *AA|A′A′*, *Aa|A′A′*, *AA|a′A′*, *Aa|A′A′*, *aa|A′A′*, *Aa|a′A′*, *aa|A′A′*, or *aa|a′A′*. Here the vertical bar ‘*|*’ denotes separation between the subgenomes and the ^*I*^ indicates homoeologous alleles. With perfect knowledge about which alleles go with each subgenome, determining the genotypes could be done completely independently. However, if separate reference genomes for the homoeologous chromosomes are not available, all reads mapping to a variable position will not be separable into reads coming from one subgenome or the other. Thus, when considering a variable site across the full set of homoeologs, we need to account for the fact that the frequency of the alternative allele may not be the same in each subgenome due to their separate evolutionary histories, even if both subgenomes are independently in Hardy Weinberg equilibrium. When we cannot separate reads, we can instead consider the full genotype of an allopolyploid with two subgenomes as being a combination of the genotypes within the subgenomes (i.e., the number of alternative alleles summed across subgenomes). Returning to the previous example, a tetraploid with two diploid subgenomes can have a full genotype of 0, *…*, 4 copies of the alternative allele, but each of these full genotypes can be found via a different combination of genotypes in the subgenomes: {0 = (0, 0); 1 = (0, 1), (1, 0); 2 = (0, 2), (2, 0), (1, 1); 3 = (1, 2), (2, 1); 4 = (2, 2)*}*. In general, for an allopolyploid that has two subgenomes with ploidy levels equal to *m*_1*i*_ and *m*_2*i*_, there are a total of (*m*_1*i*_ + 1) *×* (*m*_2*i*_ + 1) genotype combinations to consider. The probabilities of these genotypes are then determined using the allele frequencies for the alternative allele in the subgenomes.

An obvious complication of not being able to separate the sequencing reads into sets coming from each subgenome is that it makes independently estimating the allele frequencies and genotypes impossible. However, it is sometimes the case that the parental species of the allopolyploid are known, which can help with inferring genotypes by providing an outside estimate of the allele frequencies within the subgenomes. For our model, we relax this use of outside knowledge further and assume that only a single parent has been identified. Arbitrarily designating the known parent as subgenome one, we treat the allele frequencies at each locus estimated in the parental population to be known (***p**^∗^*_1_) and require only the estimation of the allele frequencies in subgenomes two (***p***_2_). We then model the full genotype in the allopolyploid as the sum of the two independent subgenomes with separate, and potentially unequal, allele frequencies. Since we assume Hardy Weinberg equilibrium within each subgenome, we can model the sum of the number of alternative alleles in the two subgenomes as a product of two binomial distributions. The log likelihood for known genotype data across individuals at all loci is then given by

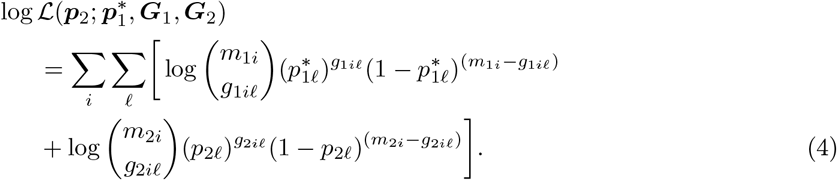

The inclusion of genotype likelihoods is done in a similar way to the autopolyploid model, only now we are summing over the values of the genotypes in both subgenomes one and two. The log likelihood for the observed sequence data given the allele frequencies in each of the subgenomes is

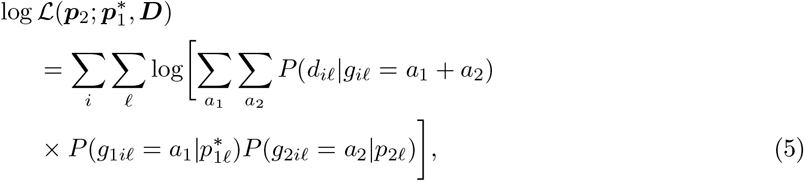

where *P* (*d_iℓ_|g_iℓ_*) is the genotype likelihood, and *P* (*g*_1*iℓ*_|p^∗^_1*ℓ*_) and *P* (*g*_2*iℓ*_|p*_2.€_*) are binomial distributions.

Because maximizing the log likelihood involves the logarithm of a double sum, we turn once again to the expectation maximization algorithm to obtain a ML estimate for the allele frequency at each locus in subgenome two (Dempster *et al*., 1977). An analytical solution for the maximization step of the EM algorithm is given by (derived in the Supplemental Text, §S1.2)

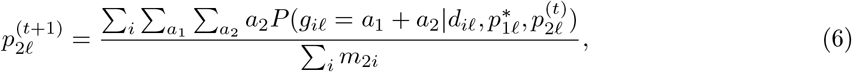

where 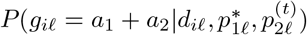 is the joint conditional probability of the genotypes in subgenomes one and two given the data and the current parameter estimates. Using these ML estimates, an empirical Bayes estimate of the genotypes within each of the subgenomes can be found using their joint posterior probability (note that subscripts *i* and *ℓ* are dropped for readability)

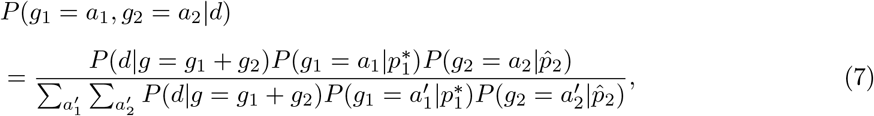

for *a*_1_ = 0, *…, m*_1*i*_ and *a*_2_ = 0, *…, m*_2*i*_.

### Other Approaches

We consider two additional approaches that use genotype priors that have been described in previous studies. The first is an implementation of the SAMtools Hardy Weinberg equilibrium prior (Li, 2011) and the second is a flat prior on genotypes that is similar to the model used by the Genome Analysis Toolkit (GATK; McKenna *et al*., 2010). Other approaches that accommodate polyploids such as the FITTETRA package in R (Voorrips *et al*., 2011) and the method of Maruki and Lynch (2017) were not considered here because they can only handle specific ploidy levels (triploids and/or tetraploids).

## Methods

Genotype likelihoods were calculated using a simplified version of the SAMtools model by using average sequencing error values at each locus, *є_ℓ_*, across reads and individuals (Li, 2011). Then for the possible values of the genotype (*a* = 0, *…, m_i_* copies of the alternative allele), the probability of the read data, *d_iℓ_* = *{t_iℓ_, r_iℓ_, є_ℓ_}* (*t_iℓ_* = total read count, *r_iℓ_* = alternative allele read count), given the genotype, *g_iℓ_*, is

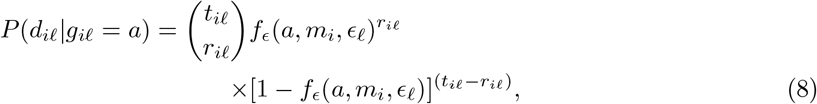

where

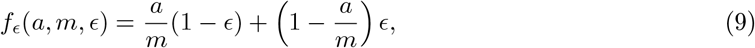

which is the probability of drawing an alternative allele weighted by the probability of a sequencing error.

### Simulations

We generated sequencing read data with mean coverage per individual, per locus equal to 2x, 5x, 10x, 20x, 30x, and 40x, simulated from a Poisson distribution for 10 000 sites. The number of individuals was set to 25, 50, or 100, and we tested ploidy levels equal to 4, 6, and 8 (4=2+2, 6=2+4, and 8=4+4 for allopolyploids). Sequencing errors were drawn from a beta distribution with parameters *α* = 1 and *β* = 200 (mean error *≈* 0.005)]. Allele frequencies were drawn from a truncated beta distribution with a minimum minor allele frequency of 5% and parameters *α* = *β* = 0.01. For the autopolyploid model, the values of the inbreeding coefficient were set to 0.1, 0.25, 0.5, 0.75, and 0.9. For the allopolyploid model, the allele frequencies simulated for subgenome one were treated as the reference panel. Genotypes were drawn according to their respective generating models (autopolyploid or allopolyploid), and the number of alternative reads for each individual at each locus was drawn from the binomial distribution in Eq.(8) given the total read count, genotype, and level of sequencing error. For each simulation, we evaluated estimation error using the root mean squared deviation (RMSD).

To compare our models with other methods, we reused these simulated data as input for the estimation of genotypes and model parameters using models that assume either Hardy Weinberg equilibrium or equal genotype probabilities (GATK-like). For the allopolyploid model, this also equates to ignoring the fact that genotypes are drawn from two independent subgenomes. Inference for the Hardy Weinberg model used the EM algorithm described in Li (2011). Genotyping based on the GATK-like model was carried out based on normalized genotype likelihoods as described in McKenna *et al*. (2010).

Comparisons for the autopolyploid model were based on the RMSD of four estimates of the inbreeding coefficient. The first of these was the estimate obtained by our ECM algorithm, which is built directly into the model. The other three estimates were calculated as a summary statistic from estimated genotypes for the three models (Supplemental Text, §S2.1). We then also compared RMSD values of the estimated genotype values for the three methods. For the allopolyploid model, direct comparisons with models that assume Hardy Weinberg or uniform genotype priors are more difficult because they do not share the assumption of two subgenomes. Therefore, we focused on the accuracy of the models to infer the full genotype by again comparing RMSD values.

### Empirical Data Analysis

#### Andropogon gerardii

We tested our autopolyploid model on an empirical data set collected in the grass species *Andropogon gerardii*. SNP data from McAllister and Miller (2016) were downloaded from Dryad as a VCF file (doi:10.5061/dryad.05qs7). The data were filtered using VCFtools v0.1.14 with the following criteria: biallelic SNPs only, no more than 50% missing data per site, one SNP per 10 000 base pair window, and a minimum sequencing depth of five reads (Danecek *et al*., 2011). The output from VCFtools was then converted to a plain text format containing the number of total reads and alternative allele reads per individual per site using a Perl script (available on GitHub). We then also removed any individuals with more than 50% missing data using an R script (available on GitHub). Since *A*. *gerardii* has two cytotypes (6N and 9N), we analyzed the hexaploid and nonaploid individuals separately and compared the estimates of the inbreeding coefficients across ploidy levels.

#### Betula pubescens and B. pendula

To test the allopolyploid model, biallelic SNP genotypes from Zohren *et al*. (2016) for the allotetraploid *Betula pubescens* and its putative diploid progenitor, *B*. *pendula*, were downloaded from Dryad (doi:10.5061/dryad.815rj). Treating the genotypes as known, we simulated read data and error values as before using Eq. (8) with beta distributed error values. We varied the level of sequencing coverage (5x, 10x, 20x) but did not alter the amount of missing data. Allele frequencies for *B*. *pendula* were estimated under the assumption of Hardy Weinberg equilibrium and disequilibrium to assess which was a better fit. These allele frequency estimates were then used as the reference panel for genotype estimation in *B*. *pubescens* using the allopolyploid model.

### Comparison with GATK

As a final comparison, we re-analyzed raw sequence data collected for *B*. *pendula* and *B*. *pubescens* using GATK v3.5.0 and our model for allopolyploids. Data for 15 individuals each of *B*. *pendula* and *B*. *pubescens* were downloaded from the European Nucleotide Archive (Project Accession ERA600270). Reads were mapped to a draft reference genome of *B*. *nana* (Dryad, doi:10.5061/dryad.815rj; Wang *et al*., 2013) using the MEM algorithm in BWA v0.7.13 with additional processing (conversion to BAM and sorting) using SAMtools v1.4.1 (Li and Durbin, 2009; Li, 2011). Read group information was added using Picard (http://broadinstitute.github.io/picard), followed by variant calling and genotype estimation using the GATK UnifiedGenotyper (*B*. *pubescens* was run with -ploidy=4; McKenna *et al*., 2010). Variant site positions in the resulting VCF files were used to extract base quality scores from the original BAM files using the SAMtools mpileup command (Li, 2011). All other data processing steps (filtering sites, finding shared variants, etc.) were conducted using Python and R scripts (available on GitHub; see Supplemental Text, §S3.2). Allele frequencies at each site were estimated in *B*. *pendula* using our implementation of the Hardy Weinberg model (run until convergence). These allele frequencies were then used as the reference panel for estimating genotypes in *B*. *pubescens* using the allopolyploid model (EM+Brent with 100 iterations). All VCF, pileup, and input/output files are publicly available on Zenodo (doi:10.5281/zenodo.825228).

#### Software and reproducibility

We have packaged our code for the EM/ECM algorithms in a C++ program called EBG, which we have included as part of a GitHub repository for this manuscript (pblischak/polyploid-genotyping; Zenodo, 10.5281/zenodo.195779). This software includes our implementations of the autopolyploid (diseq), allopolyploid (alloSNP), Hardy Weinberg (hwe), and GATK-like (gatk) models for genotyping and parameter estimation in polyploids. Each of these models also outputs the updated distribution of genotype probabilities to allow genotype uncertainty to be preserved in downstream applications. Code for the simulation study and empirical data analyses was written using a combination of the R statistical language and C++ through the use of the RCPP package (Eddelbuettel and François, 2011; Eddelbuettel, 2013; R Core Team, 2014). Figures were generated using the GGPLOT2 package in R (Wickham, 2009). Additional figure manipulations were done using Inkscape (https://inkscape.org/). All Python, Perl, R, and Bash scripts used to process data files are included on GitHub in the ‘helper-scripts/’ folder.

## Results

### Simulations

#### Autopolyploid model

Simulated read count data were generated to assess the impact of sequencing coverage and ploidy level on estimation error in autopolyploids using an expectation conditional maximization (ECM) algorithm. Convergence of the ECM algorithm depended on the number of individuals sampled, sequencing coverage, and ploidy. Each iteration of the algorithm employs Brent’s method, itself an iterative maximization algorithm, resulting in slower M-steps than the other EM algorithms we describe. However, overall convergence was reached before the maximum number of allowed iterations (1000) in all cases, with analyses typically employing between 50–100 iterations.

For the estimation of individual inbreeding coefficients (*F_i_*), Figure 1a shows the root mean squared deviation (RMSD) for estimated inbreeding coefficients for the four different estimation methods across ploidy levels and the three lowest levels of sequencing coverage (sample size of 50 individuals). Compared with the other methods that used called genotypes (diseqCG, hwe, gatk), the level of sequencing coverage and ploidy level had virtually no effect on estimation error using our model (diseq). For the other estimates, increasing sequencing coverage lowered estimation error as expected, and higher ploidy levels showed higher levels of error. However, inbreeding coefficients estimated from genotypes called from our model (diseqCG) did have lower RMSD values than the other methods, except when the inbreeding coefficient was 0.5, when the level of error was about the same. All of the methods except for Hardy Weinberg showed low levels of estimation error once the depth of sequencing reached 10x. Figures S1–S3 show the results for all simulated depths of sequencing (2x to 40x) and sample sizes (25, 50, and 100 individuals).

Our empirical Bayes approach for maximum a posteriori (MAP) genotype estimation resulted in a similar overall pattern of lower estimation error for increased sequencing coverage (Figure 1b). Interestingly, the other two methods for genotyping (gatk, hwe) showed opposing patterns of accuracy: the GATK-like model increased in accuracy with increasing levels of inbreeding but the Hardy Weinberg model had decreasing accuracy. Genotypes called by our method showed some dependence on the level of inbreeding with intermediate values having the most error. However, our method was still the most accurate across the range of inbreeding values simulated. Ploidy also had an impact on genotyping with higher ploidy levels having higher levels of estimation error. This is largely due to the fact that higher ploidy individuals have a larger number of possible values for the genotype and that the average sequencing coverage per allele (chromosome) is lower (e.g., 10x coverage in a tetraploid is on average 2.5x per allele but is 1.25x in an octoploid). Once the depth of sequencing reached 10x, the only model that still showed a higher level of error was the Hardy Weinberg model. Figures S4–S6 show the results for all simulated depths of sequencing (2x to 40x) and sample sizes (25, 50, and 100 individuals).

**Figure 1:**
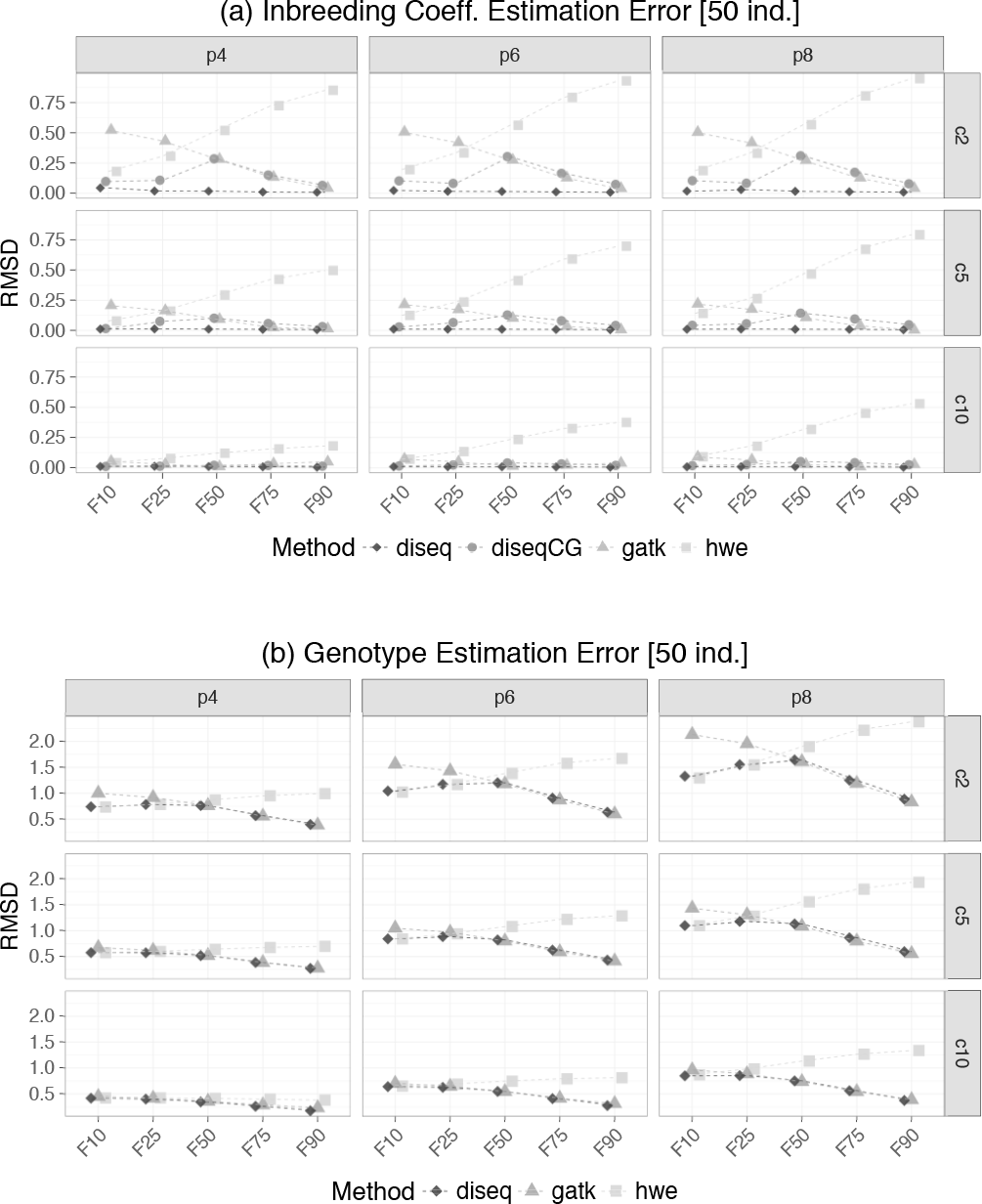
RMSD values for simulations under the autopolyploid model with inbreeding for (a) estimated inbreeding coefficients and (b) estimated genotypes. Each individual plot within (a) and (b) displays the RMSD on the y-axis and inbreeding coefficients on the x-axis. Rows correspond with the depth of sequencing coverage (2x, 5x, 10x) and the columns correspond to the ploidy level (4, 6, 8). The different estimation methods (diseq, diseqCG, gatk, hwe) are represented by different shapes within each plot. (a) The RMSD of the inbreeding coefficient estimated by our model (diseq) is consistently the lowest across all depths of sequencing coverage, ploidy level, and level of inbreeding. (b) Genotypes estimated by our model are at least as accurate as the other methods and are not as affected by high or low levels of inbreeding.

#### Allopolyploid model

Using the same general parameter settings as the simulations for the autopolyploid model (except for inbreeding), we calculated genotype likelihoods by simulating read data from genotypes generated under the model from Eq. (4). The ploidy of each subgenome was as follows: tetraploids = diploid + diploid, hexaploid = diploid + tetraploid, and octoploid = tetraploid + tetraploid. Our expectation maximization algorithm for this model was slow to converge, despite each maximization step taking less time when compared with the autopolyploid model. Analyses never reached the upper limit on the number of iterations (again 1000) but some analyses did not reach convergence until over 900 iterations had been run. To make analyses with this model more practical, we reanalyzed all simulated data sets using only 100 EM iterations followed by direct maximization of the log likelihood of the observed data given by Eq. (5) using Brent’s method (EM+Brent).

**Figure 2:**
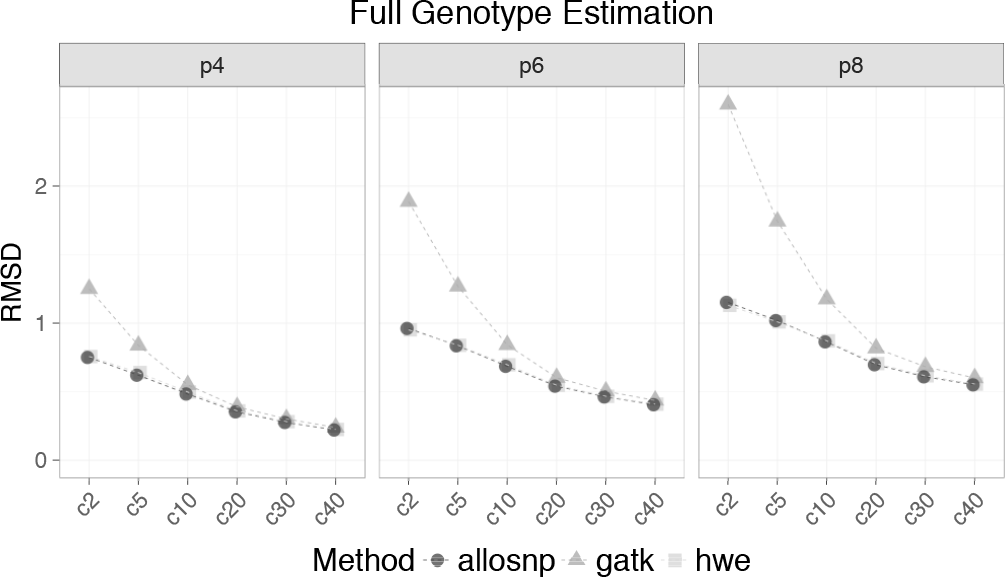
RMSD values for full genotype estimation (combined number of alternative alleles in subgenomes one and two). Sequencing coverage is on the x-axis and RMSD values are on the y-axis. Each column represents a different ploidy level and the three methods used (allosnp, gatk, hwe) are represented by different shapes. For low levels of sequencing coverage, the allosnp and hwe models have much lower levels of estimation error when compared with the gatk model. The level of sequencing coverage required for the three methods to converge in error rate depends on the ploidy level, with tetraploids needing less coverage and octoploids needing more.

Comparing our model with other genotype priors (Hardy Weinberg, GATK) only allowed us to consider the full genotype (total number of alternative alleles in subgenomes one and two) estimates from the different methods. Figure 2 shows the level of estimation error for each of the three genotyping methods for each ploidy level across all depths of sequencing coverage. For low depths of sequencing, genotyping with the GATK-like model resulted in high levels of error. As the depth of coverage increased, the three methods converged. However, this was dependent on the ploidy level: octoploids required a higher depth of sequencing for the GATK model than tetraploids or hexaploids to achieve the same level of accuracy. The Hardy Weinberg prior performed almost identically to our allopolyploid model, most likely as a result of our assuming Hardy Weinberg within the subgenomes of the allopolyploid.

We also assessed the accuracy of the model for estimating parameters based on the true values used for the simulations. Allele frequency estimates for subgenome two improved as the number of individuals and sequencing coverage were increased (Figure S7). Tetraploids showed the highest estimation error for subgenome two (diploid), followed by octoploids and hexaploids (tetraploid subgenomes), respectively. This pattern with hexaploids and octoploids is counterintuitive considering that higher ploidy levels typically result in better estimates of allele frequencies since more alleles are sampled from the population (Blischak *et al*., 2016). However, the tetraploid subgenomes in the hexaploid and octoploid individuals do not show similar levels of error as would be expected. This is likely a result of subgenome one having higher ploidy in the octoploid simulations, resulting in a larger number of possible genotype combinations and therefore higher estimation error (octoploid: 5 *×* 5 = 25 vs. hexaploid: 3 *×* 5 = 15). Figures S8 and S9 show the error in genotype estimation in subgenome one and two, respectively. Here we again observe that higher ploidy levels have higher levels of estimation error for genotypes. Overall, genotype estimates were inferred with higher error for subgenome two. This result makes sense given that we treat the allele frequencies for subgenome one as known but have to estimate them in subgenome two.

#### Empirical Data Analysis

##### Andropogon gerardii

Analyzing and filtering the data sets for hexaploid and nonaploid *A*. *gerardii* separately resulted in slightly different numbers of loci (6N: 83 individuals, 6 928 loci; 9N: 70 individuals, 6 887 loci). The average depth of sequencing coverage was 10.9x for hexaploids and 10.8x for nonaploids. Though levels of inbreeding for both cytotypes were low, nonaploids showed significantly higher levels of inbreeding than hexaploids (Figure 3a; *F*_1,151_ = 36.14, *p* = 1.3 *×* 10^*−*8^).

##### Betula pubescens and B. pendula

The data set for the species of *Betula* consisted of 130 individuals for *B*. *pubescens* and 34 individuals for *B*.*pendula* with genotype data for 49 021 loci. For *B*. *pendula*, we inferred allele frequencies and genotypes assuming Hardy Weinberg (HW), as well as using our model for individual inbreeding coefficients. The log likelihoods of the two models were very similar and most of the inbreeding coefficients were estimated to be close to 0, so we used the allele frequency estimates from the HW model as the reference panel for the allopolyploid model. After estimating the parameters of this model for *B*. *pubescens* using the EM+Brent method, we assessed the accuracy of our empirical Bayes genotype estimates by comparing them to the original data set using the root mean squared deviation (Figure 3b). The left panel shows the RMSD for each genotype value and the right panel shows a weighted measure of the RMSD that corresponds to the relative amount of error based on the frequency of that genotype in the original data set. For example, we do a poor job of estimating the genotype when the true value is 0 copies of the alternative allele, but very few of the true genotypes have that value (*∼*0.5%), so the relative contribution to the overall error is much less. In contrast, roughly 75% of the true genotypes have a value of 4 copies of the alternative allele, which is the value that we estimate the best. In addition, many of the genotypes in *B*. *pendula* were homozygous for the alternative allele (*∼*88%), so the estimates of the allele frequencies were very close to 1.0, which could have led to more error prone estimates of the genotypes in *B*. *pubescens* when using them as the reference panel.

#### Comparison with GATK

Variant calling and genotype estimation using GATK resulted in 14 931 shared SNPs between *B*. *pendula* and *B*. *pubescens* after applying the following filters: biallelic sites only, variant quality score (QUAL) greater than 30, minimum read depth (DP) per individual per site of at least five, and a maximum of five missing individuals per site. Analyzing these same sites for *B*. *pendula* using the Hardy Weinberg equilibrium model produced genotype estimates that were 99.1% identical to the estimates from GATK. Similarly for *B*. *pubescens*, genotype estimates combined from the allopolyploid subgenomes resulted in full genotype estimates that were 96.2% identical to GATK. Run times between our models and GATK are not directly comparable because it was used to identify all variants before filtering and it also performs more steps than genotyping and parameter estimation. However, it is worth noting that the analyses with our models took approximately 3.5s and 43s for the Hardy Weinberg and allopolyploid models, respectively (measured using the Unix time command).

**Figure 3:**
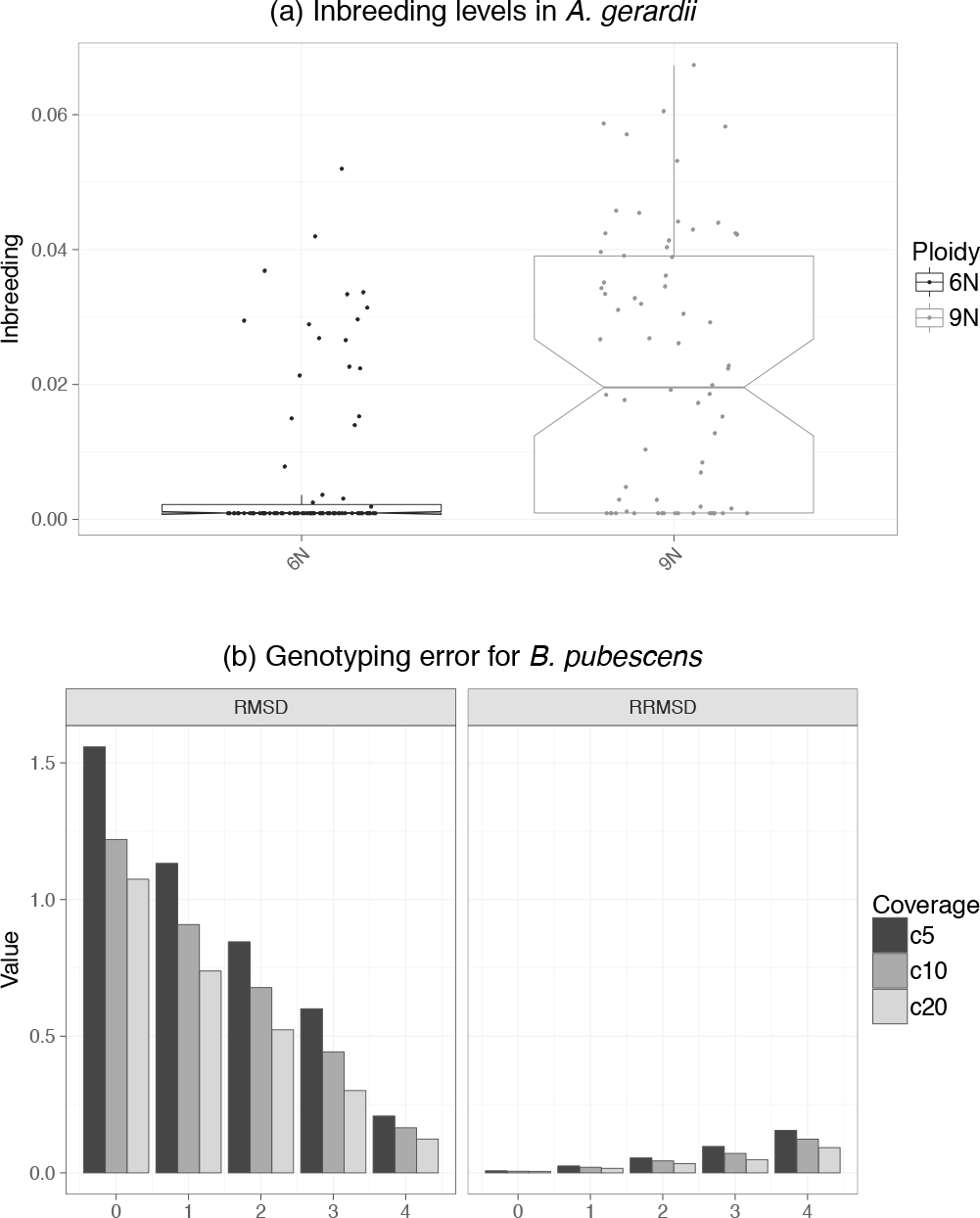
Results of empirical data analyses. (a) Levels of inbreeding in *Andropogon gerardii*. Inbreeding in the two cytotypes of *A. gerardii* is generally low, but the nonaploid (9N) samples have higher levels of inbreeding on average. (b) Genotype estimation error in *Betula pubescens*. The left panel shows the RMSD values for each of the possible full genotypes (0–4; number of alternative alleles in subgenomes one and two). The right panel shows a relative measure of the RMSD where each value is weighted by the occurrence of the particular genotype in the data set (see text for details).

## Discussion

The ability to genotype individuals in a population can be an under-appreciated task, even though it is typically the first step of any population genetic analysis. This is especially true for populations of polyploids, where genotyping is further complicated by duplicated chromosomes and their subsequent genome evolution. Until recently, genotyping polyploids using high throughput sequencing data was only possible in model organisms with reference genomes and/or subgenomes. However, more researchers have begun genotyping SNPs in both model and non-model organisms using whole genome resequencing and reduced representation methods such as restriction-site associated DNA sequencing (RADseq) and its variants (e.g., Arnold *et al*., 2015; Douglas *et al*., 2015; Cornille *et al*., 2016; Zohren *et al*., 2016). Most of these studies used already existing pieces of software to perform SNP calling and genotyping [e.g., Genome Analysis Toolkit (McKenna *et al*., 2010), UNEAK (Lu *et al*., 2012), TASSEL-GBS (Glaubitz *et al*., 2014)] but others used novel approaches for estimating genotypes (e.g., Voorrips *et al*., 2011; Zohren *et al*., 2016; Maruki and Lynch, 2017). A major caveat with these tools, however, is that many of them cannot estimate inbreeding coefficients for arbitrary ploidy levels in autopolyploids, nor can they separately estimate genotypes in the subgenomes of an allopolyploid. This is especially important considering that ignoring the independence of allopolyploid subgenomes can lead to biases in the estimation of heterozygosity when alternative alleles are fixed in the individual subgenomes (fixed heterozygosity, Cornille *et al*., 2016). In general, our models aim to incorporate more biologically realistic assumptions about how population-level factors influence the distribution of genotypes in populations of polyploids, which is critical when conducting population genetic studies in these taxa. Furthermore, our approaches use genotype likelihoods and produce updated estimates of genotype probabilities given population parameters that can be used to propagate the uncertainty in calling genotypes in polyploids to downstream analyses such as estimating heterozygosity or population differentiation, rather than relying on called genotypes.

Though our models were accurate for many of our simulations and outperformed comparable methods at low depths of sequencing coverage, it is important to consider scenarios when their assumptions are inappropriate. One concern for autopolyploids is the occurrence of double reduction, a process by which alleles in the genotype are identical by decent due to the segregation of sister chromatids to the same gamete during meiosis (Haldane, 1930). As we mentioned before, our model does not directly estimate rates of double reduction. However, because double reduction leads to identity by descent, it contributes to deviations from Hardy Weinberg that are similar to inbreeding. Therefore, our model for individual inbreeding coefficients should be able to accommodate, but not specifically estimate, double reduction.

Allopolyploids present a different set of challenges that are a result of their hybrid origins. In our model, we assume that the two subgenomes of the allopolyploid are completely independent. However, homoeologous recombination can make this assumption inappropriate. Future work that models this exchange of alleles between subgenomes will be an important extension of the model we presented here. Another potential avenue would be to develop ways to use more parental information, as well as demographic parameters to account for the amount of divergence between the allopolyploid and its parents. Models that help to identify parental taxa will also be an important contribution for future research on allopolyploids.

## Conclusions

As methods for the analysis of polyploid data continue to be developed, we are hopeful that the barriers to more widespread study of these taxa will begin to drop. The prevalence of polyploidy in plants and other groups of eukaryotes, including fish, amphibians, and fungi, make these methods fundamentally important for furthering our understanding of the impact of WGD on genetic diversity (Rogers, 1973; Otto and Whitton, 2000; Gregory and Mable, 2005; Wood *et al*., 2009). Of the main problems that complicate population genetics in polyploids, modeling allelic inheritance remains the most difficult. Overall, we believe that using genotype likelihoods when studying polyploids to overcome difficulties in determining allele copy number and for dealing with low-coverage sequencing data is a promising approach for future model development.

## Acknowledgements

The authors thank members of the Wolfe lab, B. Berger, M. Fumagalli, and three anonymous reviewers for their helpful comments on this manuscript. We also thank J. Zohren and her colleagues, C. McAllister, and A. Miller for making their data sets publicly available. Early versions of these models were presented to X. He, J. Novembre, M. Stephens, and their lab members, and we thank them for their constructive feedback and advice (especially regarding the use of EM algorithms).

## Supplementary information

Supplementary data are available online.

